# Nested Russian Doll-like Genetic Mobility Drives Rapid Dissemination of the Carbapenem Resistance Gene *bla*_KPC_

**DOI:** 10.1101/033522

**Authors:** Anna E. Sheppard, Nicole Stoesser, Daniel J. Wilson, Robert Sebra, Andrew Kasarskis, Luke W. Anson, Adam Giess, Louise J. Pankhurst, Alison Vaughan, Christopher J. Grim, Heather L. Cox, Anthony J. Yeh, the Modernising Medical Microbiology (MMM) Informatics Group, Costi D. Sifri, A. Sarah Walker, Tim E. Peto, Derrick W. Crook, AmyJ. Mathers

## Abstract

The recent widespread emergence of carbapenem resistance in *Enterobacteriaceae* is a major public health concern, as carbapenems are a therapy of last resort in this family of common bacterial pathogens. Resistance genes can mobilize via various mechanisms including conjugation and transposition, however the importance of this mobility in short-term evolution, such as within nosocomial outbreaks, is currently unknown. Using a combination of short- and long-read whole genome sequencing of 281 *bla*_KPC_-positive *Enterobacteriaceae* isolated from a single hospital over five years, we demonstrate rapid dissemination of this carbapenem resistance gene to multiple species, strains, and plasmids. Mobility of *bla*_KPC_ occurs at multiple nested genetic levels, with transmission of *bla*_KPC_ strains between individuals, frequent transfer of *bla*_KPC_ plasmids between strains/species, and frequent transposition of the *bla*_KPC_ transposon Tn*4401* between plasmids. We also identify a common insertion site for Tn*4401* within various Tn*2*-like elements, suggesting that homologous recombination between Tn*2*-like elements has enhanced the spread of Tn*4401* between different plasmid vectors. Furthermore, while short-read sequencing has known limitations for plasmid assembly, various studies have attempted to overcome this with the use of reference-based methods. We also demonstrate that as a consequence of the genetic mobility observed herein, plasmid structures can be extremely dynamic, and therefore these reference-based methods, as well as traditional partial typing methods, can produce very misleading conclusions. Overall, our findings demonstrate that non-clonal resistance gene dissemination can be extremely rapid, presenting significant challenges for public health surveillance and achieving effective control of antibiotic resistance.

**Importance:** Increasing antibiotic resistance is a major threat to human health, as highlighted by the recent emergence of multi-drug resistant “superbugs”. Here, we tracked how one important multi-drug resistance gene spread in a single hospital over five years. This revealed high levels of resistance gene mobility to multiple bacterial species, which was facilitated by various different genetic mechanisms. The mobility occurred at multiple nested genetic levels, analogous to a Russian doll set where smaller dolls may be carried along inside larger dolls. Our results challenge traditional views that drug-resistance outbreaks are due to transmission of a single pathogenic strain. Instead, outbreaks can be “gene-based”, and we must therefore focus on tracking specific resistance genes and their context rather than only specific bacteria.

## Introduction

Although antibiotic resistance genes have been identified in ancient bacterial DNA (1), much of the recent, alarming increase in pathogen antimicrobial resistance is attributable to the dissemination of resistance genes via horizontal gene transfer (HGT), in response to selection imposed by widespread antibiotic use in medicine and agriculture (2, 3). Many resistance genes are located on plasmids, which can be transferred between different bacterial strains or species, thus facilitating HGT (4). Furthermore, resistance gene mobility can be enhanced by integration into transposable elements, which are short stretches of DNA (several kilobases) that can autonomously mobilize between different genomic locations (5). However, the importance of HGT in short-term evolution is unclear, as capturing the processes in real-time is challenging, and outbreaks in health care settings are often thought to be dominated by clonal transmission (6–9).

Carbapenem resistance in *Enterobacteriaceae* has been recognized as a key threat to modern medicine (10, 11), as carbapenems often represent the therapy of last resort for serious infections (12, 13). One of the most prevalent carbapenem resistance genes is the *Klebsiella pneumoniae* carbapenemase (KPC) gene, *bla*_KPC_, first identified in 1996 and now endemic in many regions of the world (14). KPC is a beta-lactamase capable of hydrolyzing all beta-lactams, including penicillins, monobactams, cephalosporins and carbapenems (15), leaving few treatment options for infected vulnerable hospitalized patients and resulting in worse treatment outcomes (16).

Most reports of *bla*_KPC_ involve *K. pneumoniae* multi-locus sequence type (ST)258 (9, 17), which has been found globally, indicating that clonal dissemination of this resistant lineage has been an important factor in the spread of *bla*_KPC_ (9, 17–20). Nevertheless, *bla*_KPC_ has also been observed in other *K. pneumoniae* lineages, as well as other species of *Enterobacteriaceae*, suggesting that *bla*_KPC_ HGT has also played a role in resistance dissemination (21–25). As *bla*_KPC_ is often found on conjugative plasmids, some of which have been identified in multiple strains or species, this provides a likely mechanism for HGT (21, 26, 27). In addition, *bla*_KPC_ is usually present as part of the 10 kb composite Tn3-based mobile transposon Tn*4401*, which has been identified in various different plasmids, implicating Tn*4401* transposition as another mechanism contributing to *bla*_KPC_ spread (28, 29).

While Tn*4401* transposition and plasmid conjugation have been measured in the laboratory (28, 30, 31), the frequencies with which these processes occur within real-world ecosystems is not fully understood. In clinical contexts, it is often assumed that short-term evolution is dominated by clonal propagation, such that transmission chains generally involve a single pathogenic strain. However, if HGT is frequent relative to transmission (e.g. a “plasmid outbreak”), then linked patients may show variation in strain composition. If transposition is also frequent, then both host strain and resistance plasmid may show high variability within a single outbreak. As current surveillance strategies tend to focus on the host strain, it is important to establish the relevance of *bla*_KPC_ mobility within outbreak settings.

Traditional approaches for plasmid investigation, such as PCR-based replicon typing, are limited in resolution. Next-generation sequencing has been successfully applied to molecular epidemiological investigation of a number of pathogens at the host strain level, however the application and limitations of this technology for transmission chains involving HGT are relatively unexplored. Whole genome sequencing using short-read technologies (e.g. Illumina) has become cheap and accessible, but is not ideal for plasmid analysis due to *de novo* assembly limitations, as it is often not possible to accurately reconstruct the genomic context surrounding repeated sequences (21, 32). Long-read sequencing (e.g. PacBio) can largely overcome this, often providing single-contig plasmid assemblies, but it is currently prohibitively expensive for many applications. Several studies have utilised reference-based methods for plasmid assembly or inference of plasmid structures using short-read data (33, 34), however these approaches make the implicit assumption that plasmid structures are relatively stable. It will be important to understand the potential shortcomings of these assumptions in relation to mobile genetic elements, which may frequently be involved in plasmid rearrangements. Understanding when and how to successfully apply short and/or long-read sequencing technologies to molecular epidemiology tracking will be important to the field as the incidence of HGT is increasingly recognized (35).

In our institution, *bla*_KPC_ was first identified in 2007 in a patient simultaneously colonized with *bla*_KPC_-positive *K pneumoniae* and *Klebsiella oxytoca*, harbouring the *bla*_KPC_ plasmids pKPC_UVA01 and pKPC_UVA02 respectively (36, 37). Since then, we have been prospectively screening all *Enterobacteriaceae* species for *bla*_KPC_ carriage despite national guidelines which recommend screening of *Klebsiella* species and *Escherichia coli* only (38–41). Here we describe the genetic basis of non-clonal *bla*_KPC_ emergence in a single hospital setting, using a combination of short- and long-read whole genome sequencing to provide genomic characterization of 281 *Enterobacteriaceae* isolates from the first five years of this multi-species *bla*_KPC_ outbreak.

## Results

There were 204 patients infected/colonized with *bla*_KPC_-positive *Enterobacteriaceae* during the prospective sampling period, based on clinical and surveillance sampling. We performed short-read Illumina sequencing on all 294 available isolates; 13 of these were excluded due to quality issues (see Methods), leaving 281 isolates, from 182/204 (89%) patients, for analysis (Table S1). In all 281 isolates, *bla*_KPC_ was carried within a complete or partial Tn*4401* structure.

### *bla*_KPC_ is found in many different host strains, indicating frequent HGT

There were 13 different species carrying *bla*_KPC_ (Figure 1). The four most prevalent species were *Enterobacter cloacae* (96 isolates from 80 patients), *K. pneumoniae* (94 isolates from 55 patients), *Klebsiella oxytoca* (35 isolates from 20 patients), and *Citrobacter freundii* (30 isolates from 25 patients), each of which showed substantial genetic diversity. Across all species, there were a total of 62 distinct strains (>500 chromosomal SNVs; see Methods). Of these, 18 strains were identified in multiple patients, and 44 were only seen in a single patient (Figure 1), with new strains continuing to appear throughout the study period. The very recent emergence of *bla*_KPC_ on an evolutionary timescale (15) implies that each strain likely acquired *bla*_KPC_ independently, demonstrating frequent HGT between different strains and species.

**Figure 1.**
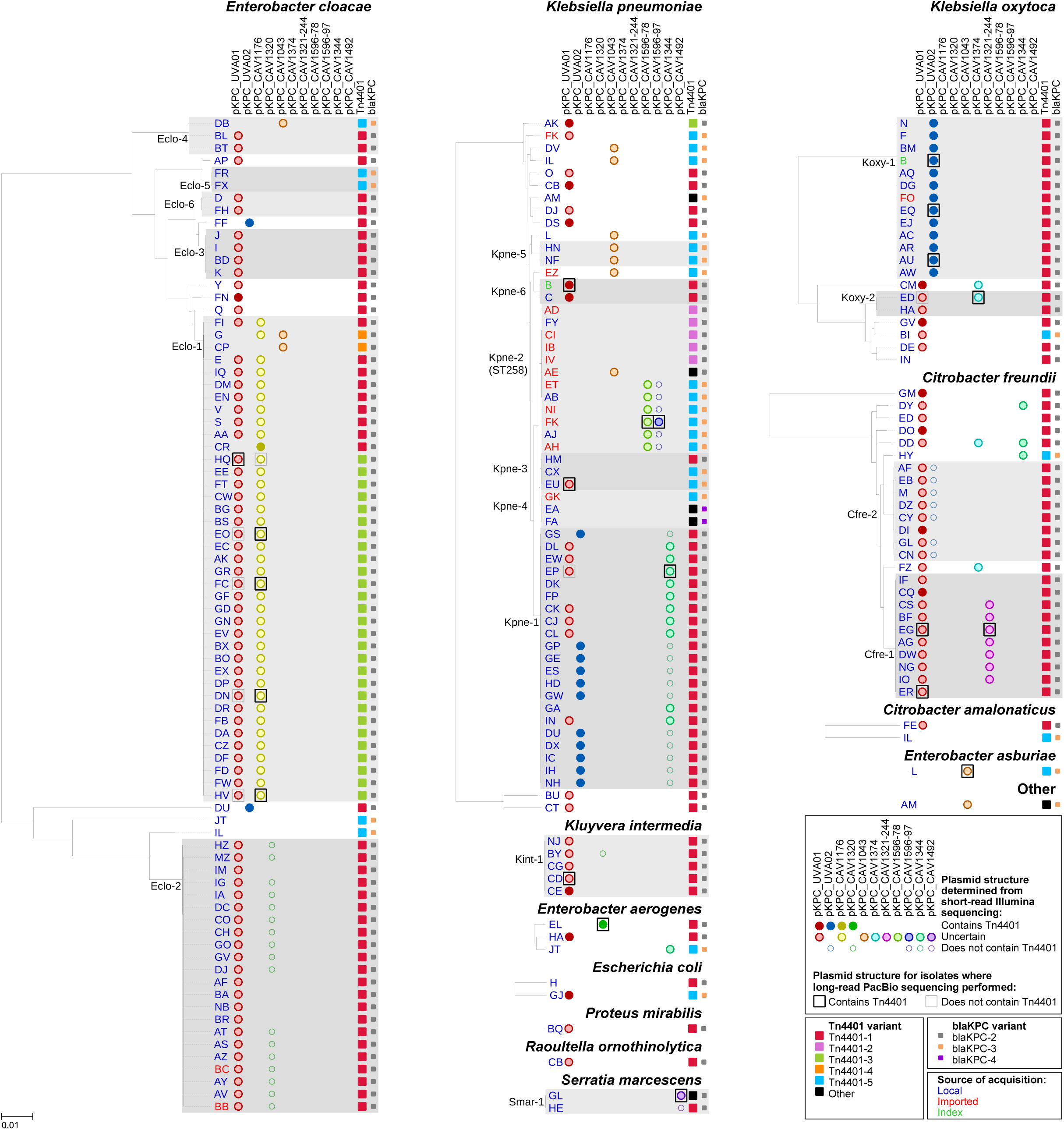
Diversity in bacterial species, strains, plasmids and Tn*4401* variants. For each species, a phylogeny was generated from mapping to a species-specific chromosomal reference, after deduplication of closely related isolates from the same patient (see Methods). Distinct strains are defined by a cutoff of ~500 SNVs (see Methods); strains found in more than one patient are indicated by grey background shading. Circles show plasmid “presence” as determined from Illumina data, with fill colour indicating uncertainty in whether the plasmid contains *bla*_KPC_· Boxes show plasmid structure determined from long-read PacBio sequencing for 17 randomly chosen isolates, as well as the previously sequenced isolates from index patient B (37). Where the PacBio-sequenced isolate was excluded from the phylogeny as a patient duplicate, plasmid structure is shown for the corresponding closely related isolate from the same patient. Tn*4401* and *bla*_KPC_ variants (Table 2) are indicated by large and small squares respectively. The likely source of *bla*_KPC_ acquisition as determined from epidemiological data is indicated by text colour.

### The *bla*_KPC_ plasmids pKPC_UVA01 and pKPC_UVA02 are widely dispersed

We hypothesised that the spread of *bla*_KPC_ could be due to conjugative transfer of the index *bla*_KPC_ plasmids, pKPC_UVA01 and pKPC_UVA02. Defining plasmid presence as ≥99% sequence identity over ≥80% of the plasmid length, 121 (66%) and 32 (18%) patients had isolates carrying pKPC_UVA01 and pKPC_UVA02, respectively, corresponding to 39 and 5 distinct strains from 10 and 4 species, respectively (Figure 1). Although the wide dispersal of these plasmids supports the plasmid-mediated outbreak hypothesis, short-read data is limited in the structural inferences it can provide when repetitive sequences are present, and for many isolates it was not possible to confirm that *bla*_KPC_ was actually co-located within pKPC_UVA01 or pKPC_UVA02 (Figure 1).

### *bla*_KPC_ is found in many different plasmids, indicating frequent Tn*4401* transposition

To further investigate *bla*_KPC_ plasmid structures, we performed long-read PacBio sequencing on 17 isolates that were chosen at random from the 281 available, yielding closed *bla*_KPC_ structures in all cases. Fifteen isolates had a single *bla*_KPC_ plasmid and two isolates had two *bla*_KPC_ plasmids, giving a total of 19 *bla*_KPC_ plasmids from the 17 isolates (Table 1). One isolate additionally had a chromosomal insertion of Tn*4401*.

**Table 1.** *bla*_KPC_-containing structures ascertained from long-read PacBio sequencing of 17 randomly chosen isolates.

| Isolate | Species | Patient | Date | *bla*<sub>KPC</sub> plasmid | Size (bp) | Group<sup>a</sup> | Within-group genetic changes<sup>b</sup> | Tn4401 variant | Flanking sequence<sup>c</sup> | Tn2-like element<sup>d</sup> |
| --- | --- | --- | --- | --- | --- | --- | --- | --- | --- | --- |
| CAV1344 | *K. pneumoniae* | EP | Dec-2010 | pKPC_CAV1344 | 176,497 | Singleton | NA | Tn4401-1 | GTTCT...GTTCT | Yes |
| CAV1392 | *K. pneumoniae* | EU | Mar-2011 | pKPC_CAV1392 | 43,621 | pKPC_UVA01 | 1 SNV | Tn4401-5 | GTTCT...GTTCT | Yes |
|  |  |  |  | NA (chromosomal) | NA | NA | NA | Tn4401-5 | AGATA...AGATA | No |
| CAV1596 | *K. pneumoniae* | FK | Apr-2012 | pKPC_CAV1596-78 | 77,801 | Singleton | NA | Tn4401-5 | GTTCT...GTTCT | Yes |
|  |  |  |  | pKPC_CAV1596-97 | 96,702 | Singleton | NA | Tn4401-5 | TATCG...TATCG | No |
| CAV1099 | *K. oxytoca* | AU | Apr-2009 | pKPC_CAV1099 | 113,105 | pKPC_UVA02 | 0 SNVs | Tn4401-1 | ATGCA...GGCCA<sup>e</sup> | No |
| CAV1335 | *K. oxytoca* | EQ | Dec-2010 | pKPC_CAV1335 | 113,105 | pKPC_UVA02 | 0 SNVs | Tn4401-1 | ATGCA...GGCCA<sup>e</sup> | No |
| CAV1374 | *K. oxytoca* | ED | Aug-2010 | pKPC_CAV1374 | 332,956 | Singleton | NA | Tn4401-1 | GTTCT...GTTCT | Yes |
| CAV1043 | *E. asburiae* | L | Mar-2008 | pKPC_CAV1043 | 59,138 | Singleton | NA | Tn4401-5 | GTTCT...GTTCT | Yes |
| CAV1176 | *E. cloacae* | DN | May-2010 | pKPC_CAV1176 | 90,452 | pKPC_CAV1176 | 0 SNVs | Tn4401-3 | GTTCT...GTTCT | Yes |
| CAV1311 | *E. cloacae* | EO | Jan-2011 | pKPC_CAV1311 | 90,452 | pKPC_CAV1176 | 0 SNVs | Tn4401-3 | GTTCT...GTTCT | Yes |
| CAV1411 | *E. cloacae* | FC | Jun-2011 | pKPC_CAV1411 | 90,452 | pKPC_CAV1176 | 1 SNV, 40 kb inversion | Tn4401-3 | GTTCT...GTTCT | Yes |
| CAV1669 | *E. cloacae* | HV | Aug-2012 | pKPC_CAV1669 | 90,452 | pKPC_CAV1176 | 40kb inversion | Tn4401-3 | GTTCT...GTTCT | Yes |
| CAV1668 | *E. cloacae* | HQ | Aug-2012 | pKPC_CAV1668 | 43,433 | pKPC_UVA01 | 1 SNV, 188 bp deletion | Tn4401-3 | GTTCT...GTTCT | Yes |
| CAV1321 | *C. freundii* | EG | Nov-2010 | pKPC_CAV1321-45 | 44,846 | pKPC_UVA01 | 1,225 bp insertion | Tn4401-1 | GTTCT...GTTCT | Yes |
|  |  |  |  | pKPC_CAV1321-244 | 243,709 | Singleton | NA | Tn4401-1 | GTTCT...GTTCT | Yes |
| CAV1741 | *C. freundii* | ER | Oct-2012 | pKPC_CAV1741 | 129,196 | pKPC_UVA01 | 14,960 bp duplication, 70,615 bp insertion | Tn4401-1<sup>f</sup> | GTTCT...GTTCT | Yes |
| CAV1151 | *K. intermedia* | CD | Sep-2009 | pKPC_CAV1151 | 43,621 | pKPC_UVA01 | 0 SNVs<sup>g</sup> | Tn4401-1 | GTTCT...GTTCT | Yes |
| CAV1320 | *E. aerogenes* | EL | Nov-2010 | pKPC_CAV1320 | 13,981 | Singleton | NA | Tn4401-1 | TTGTT...TTGTT | No |
| CAV1492 | *S. marcescens* | GL | Dec-2011 | pKPC_CAV1492 | 69,158 | Singleton | NA | Tn4401-8 | TTTTT...TTTTT | No |

<sup>a</sup> Plasmids are defined as belonging to the same group if the sequences are largely identical, allowing for a small number of substitutions and/or rearrangements that may be expected to occur within the outbreak timeframe. Different groups have very limited homology outside the Tn4401 region, indicative of independent integrations into distinct plasmid structures. 'Singleton' indicates a plasmid backbone that is distinct from all others shown
<sup>b</sup> Differences relative to the reference sequence for that plasmid group, as specified in the previous column
<sup>c</sup> Sequences immediately flanking Tn4401; generally expected to be identical due to 5 bp target site duplication during transposition (28)
<sup>d</sup> Tn4401 integrated into the *tnpA* gene of a Tn2-like element
<sup>e</sup> No evidence of target site duplication
<sup>f</sup> 2 copies
<sup>g</sup> It is noteworthy that this plasmid from CAV1151 (*K. intermedia*) is exactly identical to pKPC\_UVA01 from CAV1016 (*K. pneumoniae*), with isolation dates 2 years apart

From the analysis of Illumina data described above, 11/17 of these isolates contained pKPC_UVA01. As expected, the PacBio assemblies revealed a pKPC_UVA01-like plasmid in each of these isolates. However, only five of these pKPC_UVA01-like plasmids actually contained *bla*_KPC_ (Figure 2). The other six pKPC_UVA01-like plasmids lacked the entire Tn*4401* element, which was present on a different plasmid in these isolates. Importantly, this demonstrates that plasmid presence (as defined by Illumina sequencing) is an unreliable indicator of the mobile unit carrying *bla*_KPC_, as the “presence” of pKPC_UVA01 was misleading in 55% (6/11) of the randomly selected PacBio isolates.

**Figure 2.**
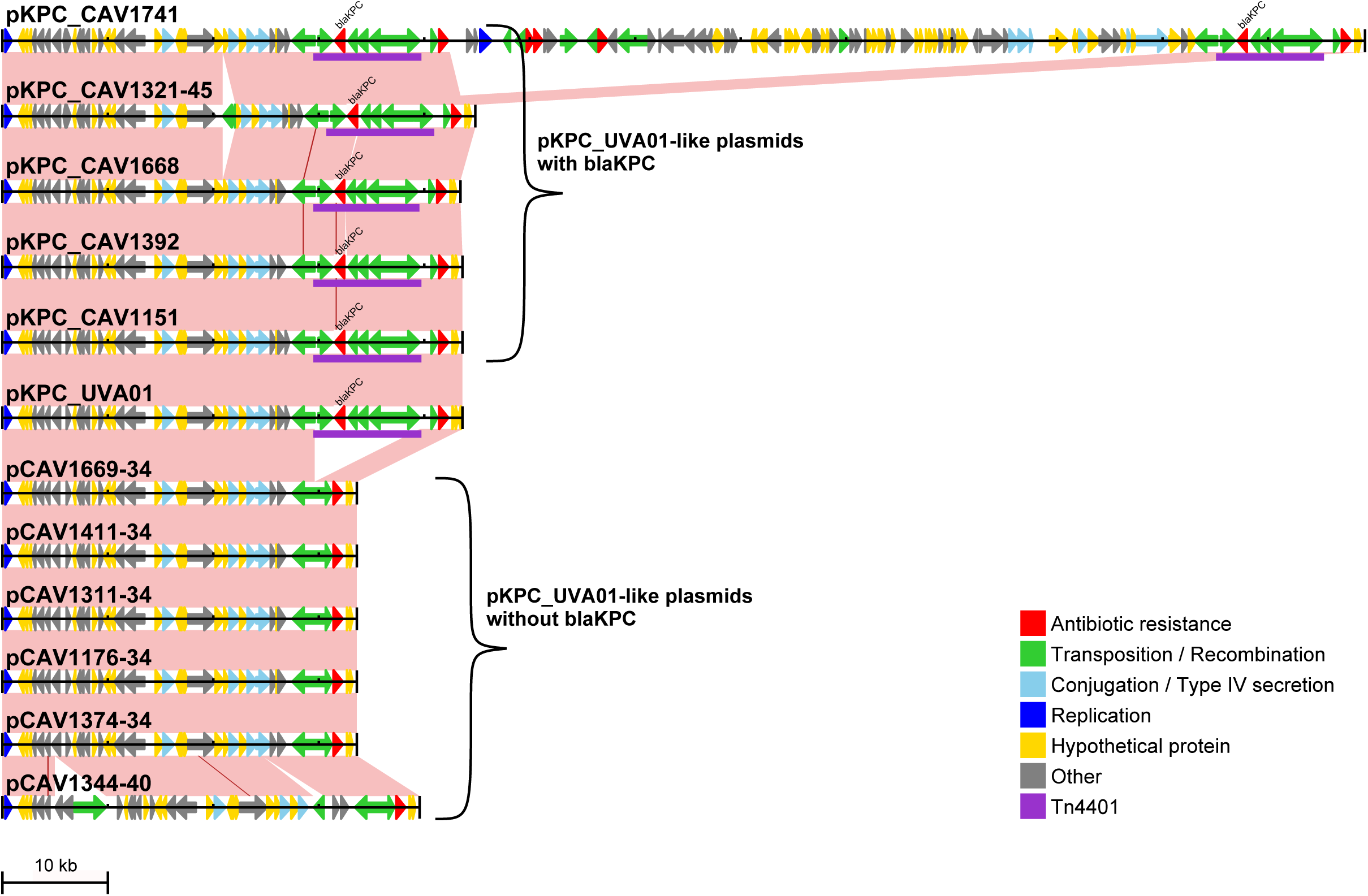
-like plasmids identified through long-read PacBio sequencing. The reference pKPC_UVA01 sequence is shown, together with all 11 pKPC_UVA01 -like plasmids identified through long-read PacBio sequencing, including the six that do not contain *bla*_KPC_. Arrows indicate predicted open reading frames; Tn*4401* is shown in purple. Pink shading indicates regions of identity between adjacent sequences, with SNVs indicated by red lines.

After accounting for multiple variants of the same plasmid backbone (e.g. the -like plasmids described above), the 19 *bla*_KPC_ plasmids identified through long-read sequencing represented 11 distinct plasmid structures (Table 1, Figure S1). These consisted of five pKPC_UVA01-like plasmids, two pKPC_UVA02-like plasmids, four pKPC_CAV1176-like plasmids, and eight *bla*_KPC_ plasmids that were each present in only a single PacBio-sequenced isolate. Using Illumina data to assess the presence of each of these 11 distinct *bla*_KPC_ plasmids across the entire set of isolates as described above revealed varied patterns of plasmid presence (Figure 1). However, in the majority of cases it was not possible to determine from Illumina data whether these plasmids contained *bla*_KPC_, so the precise details regarding distribution of *bla*_KPC_-containing plasmids across the 281 isolates remains elusive.

Taken together, these results demonstrate a great deal of *bla*_KPC_ plasmid diversity, as 11 distinct *bla*_KPC_ plasmids were identified through long-read sequencing of 17 isolates. Given that these isolates were randomly chosen, the total number of distinct *bla*_KPC_ plasmids across the entire set of 281 isolates is likely to be much greater than this. Additional Tn*4401* insertion sites were identified from the subset of isolates where flanking sequences could be adequately assembled using short-read data, further supporting this hypothesis (Table S2). Therefore, HGT of the index *bla*_KPC_ plasmids (pKPC_UVA01 and pKPC_UVA02) only partially explains *bla*_KPC_ spread, and the large number of distinct *bla*_KPC_ plasmids indicates high levels of Tn*4401* mobility.

### Tn*4401* is present within a Tn*2*-like element in many different plasmids

In 7 of the 11 distinct, fully characterised, *bla*_KPC_ plasmids, Tn*4401* was surrounded by a sequence element related to the bla_TEM-1_-containing transposon Tn*2* (Figure 3). In all cases, the insertion site of Tn*4401* within the *tnpA* gene of Tn*2* was identical, with approximately 1 kb of flanking sequence on either side of Tn*4401* showing 100% identity, but the remainder of these Tn*2*-like elements showed substantial variation. For example, while the sequence surrounding Tn*4401* in pKPC_CAV1176 was identical to the reference Tn*2*\* sequence, the Tn*2*-like element in pKPC_CAV1043 was truncated. Additionally, pKPC_CAV1344 and pKPC_CAV1596–78 contained a Tn*2* derivative, Tn *1331*, which contains the additional resistance genes *bla*_OXA-9_*, aadA1*, and *aac(6′)-lb* and has been seen as a prior site of insertion for Tn*4401* (42).

**Figure 3.**
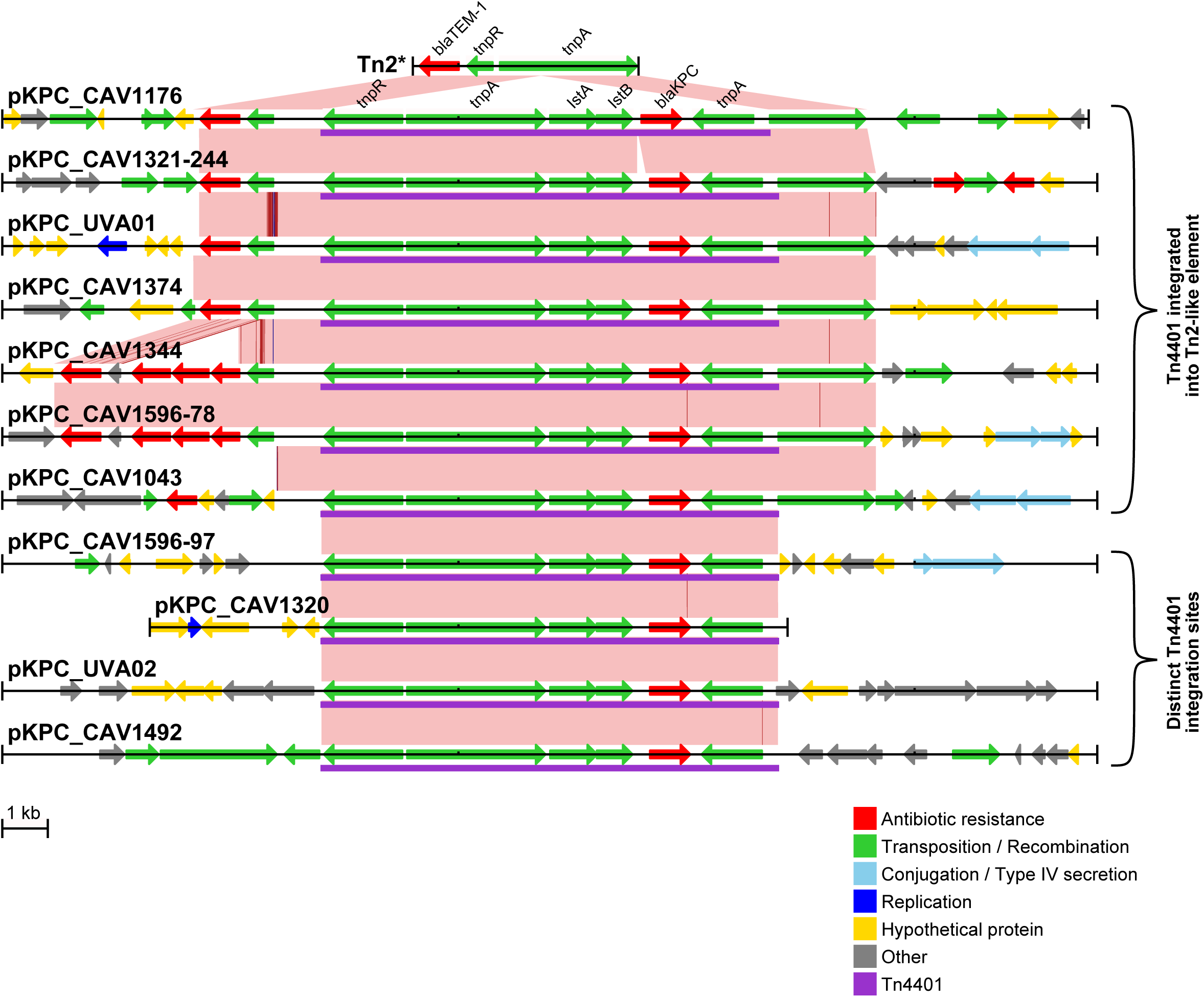
Tn*4401* is commonly integrated into a Tn*2*-like element. The Tn*4401* and surrounding region (i.e. partial plasmid sequence, except for pKPC_CAV1320) is shown for each distinct *bla*_KPC_ plasmid. Variants of the same plasmid backbone (see Table 1) are not shown. Arrows indicate predicted open reading frames; Tn*4401* is shown in purple. Pink shading indicates regions of identity between adjacent sequences, with SNVs indicated by red lines and short indels (1–2 bp) by blue lines. The top panel shows the Tn*2*\* reference sequence from AY123253 (44).

### Tn*4401* variation

There were five different structural variants of Tn*4401* (Table 2). The majority of isolates, 230/281 (82%), had the Tn*4401b* isoform, with the remaining isolates containing Tn*4401a* (n=8), a novel Tn*4401* isoform with a 188 bp deletion upstream of *bla*_KPC_ (n=39), or one of two truncated Tn*4401* structures (n=4). At the SNV level, there were seven sites that were variable within Tn*4401b*. Three of these were located within *bla*_KPC_, giving rise to three different *bla*_KPC_ alleles, *bla*_KPC-2_ (n=179), *bla*_KPC-3_ (n=44), and *bla*_KPC-4_ (n=5). All non-Tn*4401b* isolates contained *bla*_KPC_-2· Taking all structural and SNV variation into account, there were a total of 12 different Tn*4401* variants. However, most of these were very rare, with seven only found in a single patient.

**Table 2.** Tn*4401* variation.

| Tn4401 variant | Structural isoform(29) | SNVs<sup>a</sup> | bla<sub>KPC</sub> variant | Patients | Isolates | Strains |
| --- | --- | --- | --- | --- | --- | --- |
| Tn4401-1<sup>b</sup> | b | - | bla<sub>KPC-2</sub> | 121 | 176 | 42 |
| Tn4401-2 | a (del 7020-7118) | - | bla<sub>KPC-2</sub> | 5 | 8 | 1 |
| Tn4401-3 | novel (del 6919-7106) | - | bla<sub>KPC-2</sub> | 28 | 39 | 2 |
| Tn4401-4 | truncated (del 1-6654) | - | bla<sub>KPC-2</sub> | 2 | 3 | 1 |
| Tn4401-5 | b | 8015 C→T<sup>c</sup> | bla<sub>KPC-3</sub> | 22 | 40 | 19 |
| Tn4401-6 | b | 8015 C→T, 9621 T→C | bla<sub>KPC-3</sub> | 1 | 3 | 2 |
| Tn4401-7 | b | 7199 T→A, 8015 C→T, 9621 T→C | bla<sub>KPC-3</sub> | 1 | 1 | 1 |
| Tn4401-8 | b | 9663 T→C | bla<sub>KPC-2</sub> | 1 | 3 | 1 |
| Tn4401-9 | b | 7509 C→G, 7917 T→G<sup>d</sup> | bla<sub>KPC-4</sub> | 1 | 1 | 1 |
| Tn4401-10 | b | 6800 T→C, 7509 C→G, 7917 T→G | bla<sub>KPC-4</sub> | 1 | 4 | 1 |
| Tn4401-11 | b | 8015 N<sup>e</sup> | bla<sub>KPC-2</sub> / bla<sub>KPC-3</sub> | 1 | 2 | 1 |
| Tn4401-12 | truncated (del 1-6727) | 6800 N<sup>f</sup> | bla<sub>KPC-2</sub> | 1 | 1 | 1 |

<sup>a</sup> With respect to Tn4401-1, which is considered as the reference Tn4401 sequence for this study
<sup>b</sup> Tn4401-1 differs from the reference isoform b sequence in EU176013.1 by 14 SNVs, as follows: 4939 C→G, 4989 C→T, 5099 A→T, 5131 A→G, 5154 T→G, 5185 G→C, 5255 C→A, 5361 G→C, 5375 C→G, 5390 A→C, 5996 G→A, 5998 G→C, 8112 C→A, 8113 A→C
<sup>c</sup> This substitution converts bla<sub>KPC-2</sub> to bla<sub>KPC-3</sub>
<sup>d</sup> These two substitutions convert bla<sub>KPC-2</sub> to bla<sub>KPC-4</sub>
<sup>e</sup> Quality filters failed at this position due to a mixture of reads supporting C and T (i.e. Tn4401-11 actually represents a mixture of Tn4401-1 and Tn4401-5)
<sup>f</sup> Quality filters failed at this position due to a lack of reads mapped in the reverse direction. All reads mapped in the forward direction supported a reference (T) call

### *bla*_KPC_ mobility has occurred within the hospital

Based on prior healthcare exposure, *bla*_KPC_ acquisition source was classified as “imported” (likely acquisition prior to admission at our institution) for 15/182 (8%) patients and “local” (likely acquisition within our institution) for 167/182 (92%) patients (Figure 1; see Methods). Imports were more likely to be infected/colonised with *K. pneumoniae*, particularly ST258 (Table S3), consistent with previous reports of this strain being the dominant *bla*_KPC_ carrier in the US (9, 43). Thus, most host strain variation likely originated within the hospital via *bla*_KPC_ HGT. In support of this, 15/16 (94%) patients infected/colonised with multiple strains/species had shared Tn*4401* variants within the patient (Table S4), suggesting recent *bla*_KPC_ HGT. Notably, this included one patient with two different species carrying Tn*4401*–6, which is not found in any other patient.

There was also some evidence for recent within-strain Tn*4401* transposition. From the isolates that were randomly chosen for long-read sequencing, 4/17 (24%) had multiple Tn*4401* copies (Table 1). If we assume that this randomly chosen subset is representative, this extrapolates to approximately 66/281 isolates across the whole dataset. However, only 2/281 isolates had multiple Tn*4401* variants (Tn*4401*–11; Table 2), indicating that many isolates likely had multiple copies of the same Tn*4401* variant, consistent with recent Tn*4401* transposition.

Taken together, these results indicate that much of the genetic diversity observed is due to recent *bla*_KPC_ mobility, likely within the hospital ecosystem over the described five year outbreak.

### Direct patient-to-patient transmission does not explain *bla*_KPC_ acquisition

To further investigate *bla*_KPC_ acquisition source, we combined epidemiological and genetic data to trace possible transmission chains, at two different genetic levels. We considered possible transmission events where the donor and recipient were on the same ward at the same time, and carried the same host strain or Tn*4401* variant. Considering only “local” acquisitions (see above), 48/167 (29%) patients had ward contact with another patient carrying the same *bla*_KPC_-positive strain (Figure 4, top panel). A greater proportion, 106/167 (63%), of patients had ward contact with another patient carrying the same Tn*4401* variant. However, as Tn*4401*–1 is very common (66% of patients), these inferred transmissions may be spurious. With patients carrying this common variant excluded, only 15/50 (30%) had ward contact with another patient carrying the same Tn*4401* variant (Figure 4, bottom panel). Therefore, both genetic levels (strain or Tn*4401* variant) demonstrated plausible transmissions for only a minority of patients, indicating that direct patient-to-patient transmission is not the dominant mode of *bla*_KPC_ acquisition.

**Figure 4.**
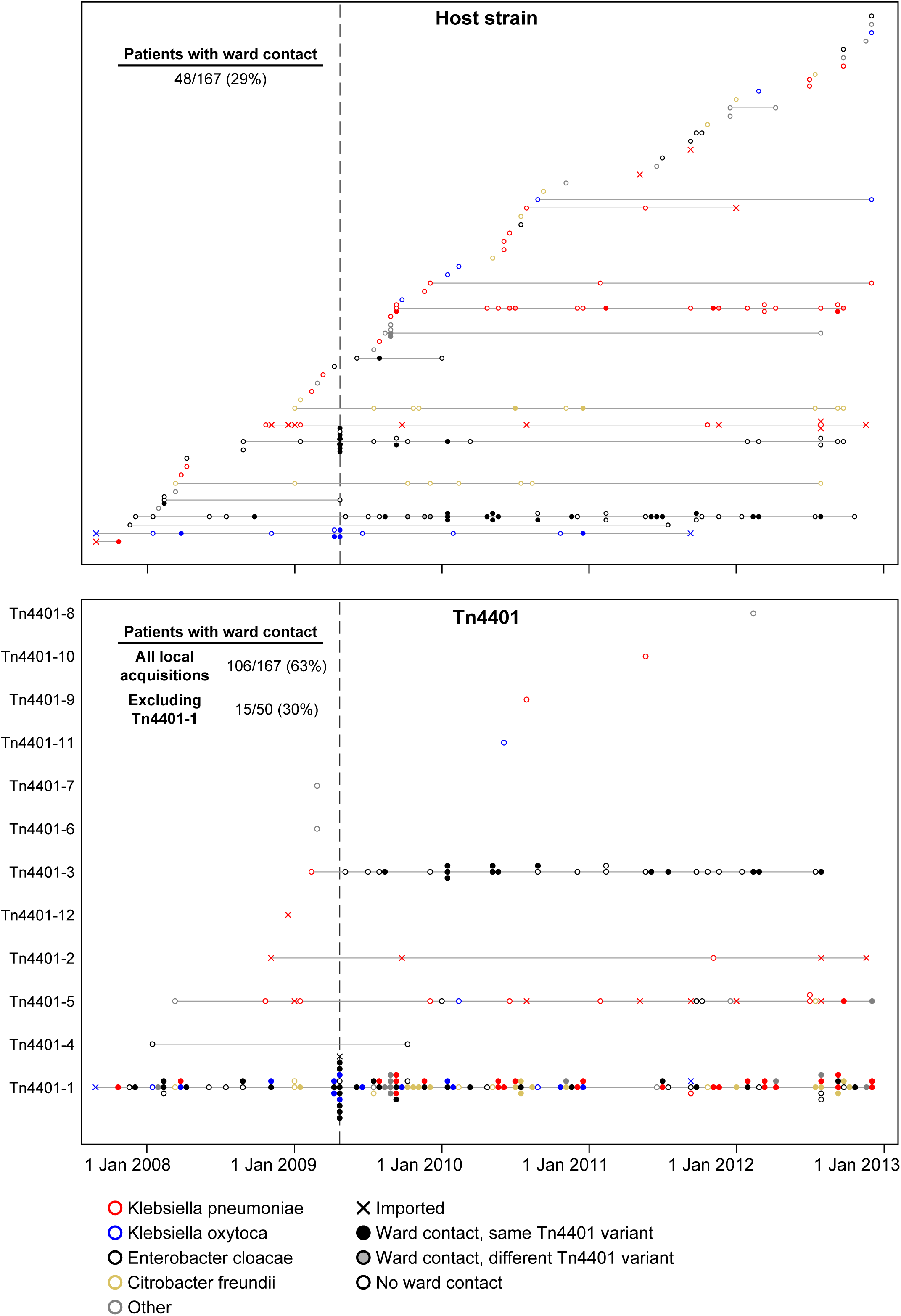
Ward contacts between patients with genetically related isolates. Each horizontal line represents a different strain (top) or Tn*4401* variant (bottom). Filled circles indicate patients that had previous ward contact with another patient on the same horizontal line (i.e. possible patient-to-patient transmission). As Tn*4401*–1 is present in two-thirds of patients, many coincidental ward contacts may be expected to occur, resulting in a substantial overestimate of transmission number. Therefore, for Tn*4401*, the total number of acquisitions explainable by direct ward contact is indicated, as well as with Tn*4401*–1 patients excluded. The vertical line indicates onset of routine patient screening.

## Discussion

Here we have demonstrated high levels of genetic diversity in KPC-producing *Enterobacteriaceae* within a single institution over five years. This diversity occurs at multiple genetic levels, revealing a complex evolutionary history of the *bla*_KPC_ gene involving many different host strains and plasmids.

In 7/11 distinct *bla*_KPC_ plasmids identified through long-read sequencing, Tn*4401* was located within a Tn*2*-like element. As these Tn*2*-like elements differed substantially from each other (Figure 3), it is unlikely that this arose via transposition of a composite Tn*4401*-Tn*2*-like structure. Instead, it suggests that Tn*4401* has been repeatedly incorporated into pre-existing Tn*2*-like elements, which are known to be widespread, and genetically divergent, in *Enterobacteriaceae* (44, 45). However, the insertion site was identical in all cases, yet Tn*4401* has been reported to have no insertion site specificity (28), suggesting that this was not facilitated by a standard transposition mechanism. Therefore, we suggest that this is most likely mediated by homologous recombination with other Tn*2*-like elements following an initial integration event, as recently suggested for another multi-drug resistance gene, *bla*_CTX-m-15_ (46). This implies that Tn*4401* mobility may have been enhanced via integration into a second, already widely dispersed, transposon. As the Tn*4401*-Tn*2*-like structure was present in the index case isolate (CAV1016, Aug 2007) we presume that the initial transposition of Tn*4401* into a Tn*2*-like element occurred prior to entry into our hospital system. In support of this, one particular Tn*2*-like element, Tn *1331*, has been previously described to contain Tn*4401* (in exactly the same position within the *tnpA* gene as described here) (21, 42, 47, 48), including one report describing a *K. pneumoniae* isolated in 2005, which predates *bla*_KPC_ in our institution (42). We are not aware of any previous reports describing Tn*4401* within a non-Tn*1331* Tn*2*-like element.

The prevalence of Tn*4401* insertions within Tn*2*-like elements also has important implications with regard to plasmid tracking. We previously published a method for arbitrary PCR to track the flanking regions around the Tn*4401* element, as well as a PCR method to assay presence of what we had wrongly assumed was a single plasmid, pKPC_UVA01. This PCR assay targeted the immediate Tn*4401* insertion site within a Tn*2*-like element (49), which we have here demonstrated is present in many different plasmids, highlighting that PCR assays, and indeed any partial typing methods, need to be interpreted with a great deal of caution. We were further mislead by the analysis of short-read whole genome sequencing data, which indicated presence of pKPC_UVA01 in the majority of isolates. Taken together it was tempting to conclude that horizontal transfer of pKPC_UVA01 was responsible for the vast majority of *bla*_KPC_ carriage in our institution. However, long-read sequencing refuted this, revealing a far more complex picture.

More generally, this highlights certain limitations for plasmid reconstruction from short-read data. To illustrate by way of example, there were five isolates where long-read sequencing revealed pKPC_UVA01-like plasmids that were identical to the reference pKPC_UVA01 sequence apart from the absence of Tn*4401* and associated 5 bp target site duplication (Figure 2). We presume that in these lineages, *bla*_KPC_ may have been initially acquired via HGT of pKPC_UVA01, with subsequent homologous recombination transferring Tn*4401* from pKPC_UVA01 to a different plasmid containing a Tn*2*-like element. In each of these five isolates, multiple Tn*2*-like elements are present, which have 100% sequence identity over approximately 1 kb on either side of the Tn*4401* insertion site. As this is longer than the fragment length used for paired-end sequencing, it is not possible to resolve the plasmid context of *bla*_KPC_ using short-read data. Importantly, any reference-based method for plasmid reconstruction (e.g. in this case using the pKPC_UVA01 reference sequence to infer presence of the plasmid in each isolate) is liable to produce misleading results. More generally, it is exactly the repetitive regions that cannot be resolved using short-read data that could be expected to be involved in plasmid rearrangements, either through homologous recombination as suggested here, or by virtue of the fact that transposable elements are often present in multiple copies. Therefore, having short-read data that is consistent with a known plasmid structure, even within the same outbreak, should not be sufficient to conclude that that structure is present, if the data is also consistent with an alternative structure. As several recent studies have utilised reference-based approaches for plasmid assembly / inference (33, 34), our results indicate that any such methods should be interpreted with extreme caution.

Across the *bla*_KPC_-positive patients, there was large variation in both host strains and *bla*_KPC_ plasmids, with Tn*4401* being the largest genetic unit that was consistently present. Therefore, surveillance strategies aimed at tracking individual strains or plasmids could be misleading, and it may be more appropriate to focus on Tn*4401*. However, we found limited variation within the transposon, as Tn*4401* sequences from 121/182 (66%) patients were identical to the index case (Table 2). This lack of variation implies that even the highest resolution genetic methods may be insufficient for determining specific transmission routes. Even so, we have demonstrated that only a minority of *bla*_KPC_ acquisition events can be explained by direct patient-to-patient transmission. Future studies should therefore contemporaneously investigate the possible involvement of unsampled reservoirs (e.g. environmental or silent colonization by additional carriers).

There several limitations to this study. Because of the cost and effort involved in long-read sequencing, we were only able to resolve a minority of *bla*_KPC_-plasmids. This means that although we have a compelling indicator of the diversity created by mobile genetic elements within a single hospital over a five year period, we are limited in the ability to genetically resolve pathways of *bla*_KPC_ mobility between host strains and plasmid vectors, even within a single patient. We also speculate about the effect of Tn*4401* insertion into Tn*2*-like elements, but future *in vitro* studies could be used to illuminate the effect of this composite structure on Tn*4401* mobility.

In conclusion, our detailed genetic analysis of the evolutionary events occurring in the early stages of antimicrobial resistance gene emergence in a single institution identifies several distinct processes occurring at high frequency (Figure 5). First, the presence of shared *bla*_KPC_**-** containing strains in different patients reflects traditional (clonal) outbreak models. Second, *bla*_KPC_ mobility between strains/species is facilitated by promiscuous *bla*_KPC_ plasmids such as pKPC_UVA01. Third, *bla*_KPC_ transfer between plasmids is likely enhanced by homologous recombination between Tn*2*-like elements, facilitating the movement of Tn*4401* from one plasmid to another. Finally, *bla*_KPC_ mobility is also enabled by standard Tn*4401* transposition. Rather than a single process dominating, resistance dissemination is driven by a combination of these factors, resulting in a high level of diversity in KPC-producing *Enterobacteriaceae*, at multiple genetic levels. As *bla*_KPC_ prevalence continues to increase, so will this genetic diversity, inevitably resulting in a wider variety of more pathogenic strains carrying *bla*_KPC_*·*

**Figure 5.**
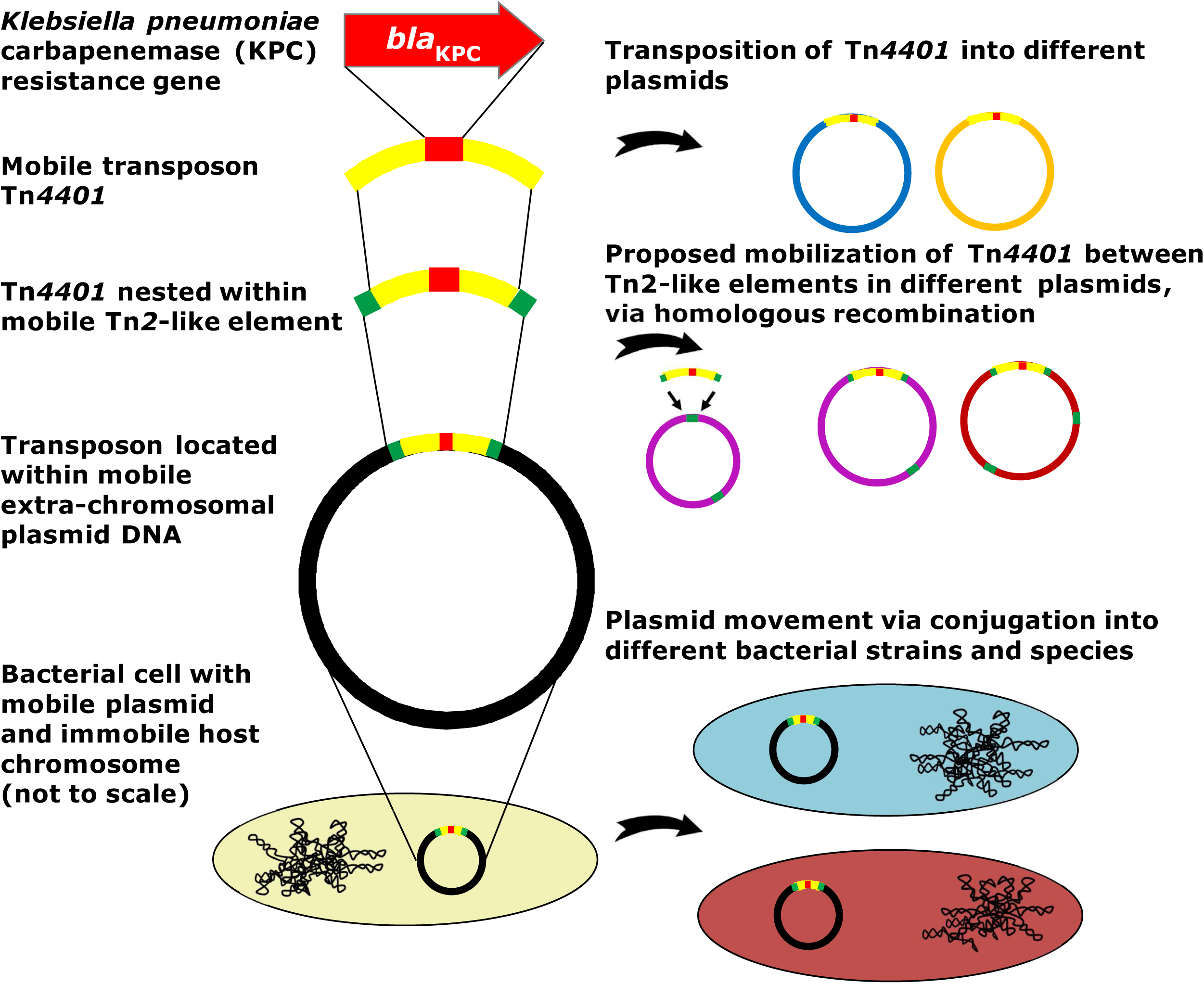
*bla*_KPC_ spreads at multiple genetic levels, resulting in a high level of diversity in *bla*_KPC_-positive *Enterobacteriaceae*.

Our results indicate that the current standard practice of only screening specific species for *bla*_KPC_ carriage is likely to hamper surveillance efforts by grossly underestimating true prevalence. Instead of the traditional view of an outbreak involving a single pathogenic strain, we propose that for KPC-producing *Enterobacteriaceae*, and possibly more generally, we should instead adopt the view of a “gene-based outbreak”, with surveillance strategies tracking the resistance gene itself rather than a specific host strain.

## Methods

### Isolate collection and Illumina sequencing

A subset of *K. pneumoniae* isolates, with corresponding sequence data, have been previously described (37). Isolate collection, *de novo* assembly, mapping and variant calling were performed as previously described (37), however here we used species-specific references for mapping (Table S5). Illumina sequencing was also performed as previously described (37), with some exceptions (see Supplementary Methods). In total, 281 isolates from 182 patients were available for analysis; exclusion criteria for additional isolates is described in Supplementary Methods.

### Species classification

Species classification was performed using microbiological and sequenced-based methods (see Supplementary Methods for details).

### Phylogenetic analysis and strain classification

There were 52 patients with multiple isolates of the same species. One of these (patient FK) involved two strains of *K. pneumoniae* that were highly divergent from each other (>20,000 chromosomal SNVs), clearly representing a separate *bla*_KPC_ acquisition by each strain. Excluding this divergent strain pair, the remaining cases had SNV differences ranging from 0 to 60 (median 2 SNVs). As these could plausibly represent clonal evolution within the patient, we conservatively chose to include only a single representative (the earliest isolate) for phylogenetic reconstruction, in order to avoid artificially inflating genetic clusters due to repeated patient sampling. Phylogenetic analysis was then performed separately for each species using PhyML (50) (see Supplementary Methods). Chromosomally distinct strains were defined by partitioning each phylogeny with a cutoff of ~500 SNVs (see Supplementary Methods). Based on the molecular clock of *Enterobacteriaceae* (1–20 SNVs/chromosome/year) (6, 37, 51), we can be relatively confident that isolates belonging to distinct strains will not have a shared ancestor within the timeframe of *bla*_KPC_ dispersal, and the number of distinct strains thus provides a conservative estimate of the number of distinct *bla*_KPC_ acquisition events.

### Long-read PacBio sequencing

For long-read sequencing, 17 isolates were randomly chosen from the entire set of sequenced isolates (i.e. including patient duplicates). Long-read sequencing and initial *de novo* assembly were performed as previously described (37). Refinement of assemblies and closure of plasmid/chromosomal sequences was performed as described in Supplementary Methods.

Since the isolates for PacBio sequencing were randomly chosen from the set of all Illumina sequenced isolates, some of them represented within-patient strain duplicates (see previous section on phylogenetic analysis), and were therefore not included in phylogenetic reconstruction. For display purposes (in Figure 1), the *bla*_KPC_ structure(s) determined from long-read PacBio sequencing for each of these isolates is shown alongside the representative isolate of the same strain from the same patient. In all cases, the representative isolate has the same short-read plasmid profile and Tn*4401* variant as the PacBio sequenced isolate.

### Plasmid presence / absence classification

The index *bla*_KPC_ plasmids pKPC_UVA01 and pKPC_UVA02, together with the additional nine distinct *bla*_KPC_ plasmids identified though long-read PacBio sequencing, were used as references to determine plasmid presence profiles for each isolate based on the Illumina data. Plasmid presence was defined as ≥99% sequence identity over ≥80% of the length of the reference sequence, as determined by BLASTn comparisons between each isolate’s *de novo* assembly and the reference plasmid. The high identity cutoff was chosen to reduce false positives from sequences that are only distantly related (and therefore unlikely to have a common ancestor within the timeframe of the outbreak), while the more permissive length cutoff allows for some rearrangement. It should be noted that the method does not take any account of structural continuity.

### Analysis of Tn*4401* flanking sequences

Where a plasmid was classified as being present in a particular isolate, it was not always certain to contain Tn*4401*. The plasmid presence classification was further refined as: “containing Tn*4401”* if the isolate’s *de novo* assembly supported Tn*4401* being present within the expected sequence context of that plasmid, “not containing Tn*4401”* if the plasmid was assembled without Tn*4401*, or “uncertain” if structure could not be determined from the *de novo* assembly. The identification of novel Tn*4401* insertion sites was also based on the *de novo* assemblies. These methods are described in detail in Supplementary Methods.

### Variation in Tn*4401*

Tn*4401* isoform classification was performed by comparing each isolate’s *de novo* assembly with the previously described isoform b reference sequence from EU176013.1 (29) using BLASTn, to identify structural variation. SNV variation was determined by mapping to a reference consisting of pKPC_UVA01 plus a species-specific chromosome as described above, followed by extraction of the Tn*4401* region. Variation is reported for all sites where at least one isolate had a non-reference call, including any ambiguity at that site in other isolates. Ambiguity at non-variable sites is not reported, which may result in an underestimate of true variation. However, any resulting underestimation is likely to be very minor, as the proportion of called sites, excluding deleted regions described above, was >96% for all isolates.

### Epidemiological classification

For epidemiologic analysis, patients were assigned a one or two letter code for de-identification. Routine peri-rectal surveillance cultures for silent colonization began in April 2009 (38, 40). Patients were classified as “imported” if they did not have any prior admission to University of Virginia Medical Center/Long-term Acute Care Hospital (UVaMC) and either had a *bla*_KPC_-positive *Enterobacteriaceae* isolated within 48 hours of admission, or had a carbapenem-resistant *Enterobacteriaceae* culture before transfer to UVaMC with a subsequent isolate at UVaMC confirmed as *bla*_KPC_ PCR positive. The index case was also classified as imported. For the remaining patients, the source of *bla*_KPC_ acquisition was classified as “local”. The 48h cutoff is arbitrary and may result in some misclassification if patients either acquire *bla*_KPC_ within the first 48h of admission, or if *bla*_KPC_ carriage/infection remains undetected for >48h, however this is expected to be minimal (see Supplementary Methods). Charts and patient contacts were reviewed using bed tracing data and the electronic medical record. The study was approved by the University of Virginia Institutional Review Board (protocol # 13558).

### Transmission analysis

Possible patient-to-patient transmission events were determined on the basis of having overlapping stays on the same ward, as well as genetically-related *bla*_KPC_ isolates. The analysis was performed separately for two different levels of genetic relatedness (strain or Tn*4401* variant). This is described in detail in Supplementary Methods.

## Acknowledgements

We thank the UVaMC Clinical Microbiology staff for collection of study isolates and the UVaMC Infection Prevention and Control staff for assistance with patient tracking.

### Modernising Medical Microbiology (MMM) informatics group

Jim Davies, Charles Crichton, Milind Acharya, Carlos del Ojo Elias

### Additional Information

Sequence data has been deposited with the National Centre for Biotechnology Information (NCBI) under BioProject PRJNA246471.

## Supplementary Legends

**Figure S1. Distinct *bla*_KPC_ plasmids identified through long-read PacBio sequencing**. Variants of the same plasmid backbone (see Table 1) are not shown. Arrows indicate predicted open reading frames; Tn*4401* is shown in purple.

**Table S1. Details of sequenced isolates.**

**Table S2. Additional Tn*4401* insertion sites ascertained from short-read Illumina data**

**Table S3. Association of importation status with *K. pneumoniae* and the epidemic *bla*_KPC_ *K. pneumoniae* strain ST258**

**Table S4. Patients with multiple *bla*_KPC_-positive strains or species**

**Table S5. Chromosomal references used for mapping**

## References

1. D’Costa VM, King CE, Kalan L, Morar M, Sung WW, Schwarz C, Froese D, Zazula G, Calmels F, Debruyne R, Golding GB, Poinar HN, Wright GD. 2011. Antibiotic resistance is ancient. Nature 477: 457–461.

2. Datta N, Hughes VM. 1983. Plasmids of the same Inc groups in Enterobacteria before and after the medical use of antibiotics. Nature 306: 616–617.

3. Hughes VM, Datta N. 1983. Conjugative plasmids in bacteria of the ‘pre-antibiotic’ era. Nature 302: 725–726.

4. Carattoli A. 2013. Plasmids and the spread of resistance. Int J Med Microbiol 303: 298–304.

5. Partridge SR. 2011. Analysis of antibiotic resistance regions in Gram-negative bacteria. FEMS Microbiol Rev 35: 820–855.

6. Stoesser N, Giess A, Batty EM, Sheppard AE, Walker AS, Wilson DJ, Didelot X, Bashir A, Sebra R, Kasarskis A, Sthapit B, Shakya M, Kelly D, Pollard AJ, Peto TE, Crook DW, Donnelly P, Thorson S, Amatya P, Joshi S. 2014. Genome Sequencing of an Extended Series of NDM-Producing Klebsiella pneumoniae Isolates from Neonatal Infections in a Nepali Hospital Characterizes the Extent of Community-versus Hospital-Associated Transmission in an Endemic Setting. Antimicrob Agents Chemother 58: 7347–7357.

7. Poirel L, Savov E, Nazli A, Trifonova A, Todorova I, Gergova I, Nordmann P. 2014. Outbreak caused by NDM-1- and RmtB-producing Escherichia coli in Bulgaria. Antimicrob Agents Chemother 58: 2472–2474.

8. Garza-Ramos U, Barrios H, Reyna-Flores F, Sánchez-Pérez A, Tamayo-Legorreta E, Ibarra-Pacheco A, Salazar-Salinas J, Núñez-Ceballos R, Silva-Sanchez J. 2014. Characteristics of KPC-2-producing Klebsiella pneumoniae (ST258) clinical isolates from outbreaks in 2 Mexican medical centers. Diagn Microbiol Infect Dis 79: 483–485.

9. Kitchel B, Rasheed JK, Patel JB, Srinivasan A, Navon-Venezia S, Carmeli Y, Brolund A, Giske CG. 2009. Molecular epidemiology of KPC-producing *Klebsiella pneumoniae* isolates in the United States: clonal expansion of multilocus sequence type 258. Antimicrob Agents Chemother 53: 3365–3370.

10. World Health Organization. 2014. Antimicrobial Resistance A Global Report on Surviellence. World Health Organization, Press W, 20 Avenue Appia, 1211 Geneva 27, Switzerland.

11. Centers for Disease Control and Prevention (CDC). 2013. Antibiotic resistance threats in the United States. (CDC) CfDCaP, Atlanta, GA USA.

12. Patel G, Huprikar S, Factor SH, Jenkins SG, Calfee DP. 2008. Outcomes of carbapenem-resistant Klebsiella pneumoniae infection and the impact of antimicrobial and adjunctive therapies. Infect Control Hosp Epidemiol 29: 1099–1106.

13. Falagas ME, Tansarli GS, Karageorgopoulos DE, Vardakas KZ. 2014. Deaths attributable to carbapenem-resistant enterobacteriaceae infections. Emerg Infect Dis 20: 1170–1175.

14. Munoz-Price LS, Poirel L, Bonomo RA, Schwaber MJ, Daikos GL, Cormican M, Cornaglia G, Garau J, Gniadkowski M, Hayden MK, Kumarasamy K, Livermore DM, Maya JJ, Nordmann P, Patel JB, Paterson DL, Pitout J, Villegas MV, Wang H, Woodford N, Quinn JP. 2013. Clinical epidemiology of the global expansion of Klebsiella pneumoniae carbapenemases. Lancet Infect Dis 13: 785–796.

15. Yigit H, Queenan AM, Anderson GJ, Domenech-Sanchez A, Biddle JW, Steward CD, Alberti S, Bush K, Tenover FC. 2001. Novel carbapenem-hydrolyzing beta-lactamase, KPC-1, from a carbapenem-resistant strain of Klebsiella pneumoniae. Antimicrob Agents Chemother 45: 1151–1161.

16. Tzouvelekis LS, Miriagou V, Kotsakis SD, Spyridopoulou K, Athanasiou E, Karagouni E, Tzelepi E, Daikos GL. 2013. KPC-Producing, Multi-Drug Resistant Klebsiella pneumoniae ST258 as a Typical Opportunistic Pathogen. Antimicrob Agents Chemother.

17. Leavitt A, Carmeli Y, Chmelnitsky I, Goren MG, Ofek I, Navon-Venezia S. 2010. Molecular epidemiology, sequence types, and plasmid analyses of KPC-producing Klebsiella pneumoniae strains in Israel. Antimicrob Agents Chemother 54: 3002–3006.

18. Adler A, Hussein O, Ben-David D, Masarwa S, Navon-Venezia S, Schwaber MJ, Carmeli Y, Group obotP-A-CHC-REW. 2014. Persistence of Klebsiella pneumoniae ST258 as the predominant clone of carbapenemase-producing Enterobacteriaceae in post-acute-care hospitals in Israel, 2008–13. J Antimicrob Chemother.

19. Deleo FR, Chen L, Porcella SF, Martens CA, Kobayashi SD, Porter AR, Chavda KD, Jacobs MR, Mathema B, Olsen RJ, Bonomo RA, Musser JM, Kreiswirth BN. 2014. Molecular dissection of the evolution of carbapenem-resistant multilocus sequence type 258 *Klebsiella pneumoniae*. Proc Natl Acad Sci U S A 111: 4988–4993.

20. Chen L, Mathema B, Chavda KD, DeLeo FR, Bonomo RA, Kreiswirth BN. 2014. Carbapenemase-producing Klebsiella pneumoniae: molecular and genetic decoding. Trends Microbiol 22: 686–696.

21. Conlan S, Thomas PJ, Deming C, Park M, Lau AF, Dekker JP, Snitkin ES, Clark TA, Luong K, Song Y, Tsai YC, Boitano M, Dayal J, Brooks SY, Schmidt B, Young AC, Thomas JW, Bouffard GG, Blakesley RW, Mullikin JC, Korlach J, Henderson DK, Frank KM, Palmore TN, Segre JA, Program NCS. 2014. Single-molecule sequencing to track plasmid diversity of hospital-associated carbapenemase-producing Enterobacteriaceae. Sci Transl Med 6:254ra126.

22. Ruiz-Garbajosa P, Curiao T, Tato M, Gijon D, Pintado V, Valverde A, Baquero F, Morosini MI, Coque TM, Canton R. 2013. Multiclonal dispersal of KPC genes following the emergence of non-ST258 KPC-producing Klebsiella pneumoniae clones in Madrid, Spain. J Antimicrob Chemother.

23. Ocampo AM, Chen L, Cienfuegos AV, Roncancio G, Chavda KD, Kreiswirth BN, Jiménez JN. 2015. High frequency of non-CG258 clones of carbapenem-resistant *Klebsiella pneumoniae* with distinct clinical characteristics: A two-year surveillance in five Colombian tertiary care hospitals. Antimicrob Agents Chemother.

24. Tavares CP, Pereira PS, Marques EeA, Faria C, de Souza MaP, de Almeida R, Alves CeF, Asensi MD, Carvalho-Assef AP. 2015. Molecular epidemiology of KPC-2-producing Enterobacteriaceae (*non-Klebsiella pneumoniae*) isolated from Brazil. Diagn Microbiol Infect Dis 82: 326–330.

25. Bonura C, Giuffrè M, Aleo A, Fasciana T, Di Bernardo F, Stampone T, Giammanco A, Palma DM, Mammina C, Group M-GW. 2015. An Update of the Evolving Epidemic of *bla*_KPC_ Carrying *Klebsiella pneumoniae* in Sicily, Italy, 2014: Emergence of Multiple Non-ST258 Clones. PLoS One 10:e0132936.

26. Chen L, Chavda KD, Melano RG, Jacobs MR, Koll B, Hong T, Rojtman AD, Levi MH, Bonomo RA, Kreiswirth BN. 2014. Comparative genomic analysis of KPC-encoding pKpQIL-like plasmids and their distribution in New Jersey and New York Hospitals. Antimicrob Agents Chemother 58: 2871–2877.

27. Tijet N, Muller MP, Matukas LM, Khan A, Patel SN, Melano RG. 2015. Lateral dissemination and inter-patient transmission of *bla*_KPC-3_: role of a conjugative plasmid in spreading carbapenem resistance. J Antimicrob Chemother.

28. Cuzon G, Naas T, Nordmann P. 2011. Functional characterization of Tn*4401*, a Tn3-based transposon involved in *bla*_KPC_ gene mobilization. Antimicrob Agents Chemother 55: 5370–5373.

29. Naas T, Cuzon G, Villegas MV, Lartigue MF, Quinn JP, Nordmann P. 2008. Genetic structures at the origin of acquisition of the beta-lactamase bla KPC gene. Antimicrob Agents Chemother 52: 1257–1263.

30. Siu LK, Lin JC, Gomez E, Eng R, Chiang T. 2012. Virulence and plasmid transferability of KPC Klebsiella pneumoniae at the Veterans Affairs Healthcare System of New Jersey. Microb Drug Resist 18: 380–384.

31. Cai JC, Zhou HW, Zhang R, Chen GX. 2008. Emergence of Serratia marcescens, Klebsiella pneumoniae, and Escherichia coli Isolates possessing the plasmid-mediated carbapenem-hydrolyzing beta-lactamase KPC-2 in intensive care units of a Chinese hospital. Antimicrob Agents Chemother 52: 2014–2018.

32. Brouwer MS, Tagg KA, Mevius DJ, Iredell JR, Bossers A, Smith HE, Partridge SR. 2015. Incl shufflons: Assembly issues in the next-generation sequencing era. Plasmid 80: 111–117.

33. Lanza VF, de Toro M, Garcillän-Barcia MP, Mora A, Blanco J, Coque TM, de la Cruz F. 2014. Plasmid flux in *Escherichia coli* ST131 sublineages, analyzed by plasmid constellation network (PLACNET), a new method for plasmid reconstruction from whole genome sequences. PLoS Genet 10:e1004766.

34. Pecora ND, Li N, Allard M, Li C, Albano E, Delaney M, Dubois A, Onderdonk AB, Bry L. 2015. Genomically Informed Surveillance for Carbapenem-Resistant Enterobacteriaceae in a Health Care System. MBio 6:e01030.

35. Partridge SR. 2015. Resistance mechanisms in Enterobacteriaceae. Pathology 47: 276–284.

36. Mathers AJ, Cox HL, Bonatti H, Kitchel B, Brassinga AK, Wispelwey B, Sawyer RG, Pruett TL, Hazen KC, Patel JB, Sifri CD. 2009. Fatal cross infection by carbapenem-resistant Klebsiella in two liver transplant recipients. Transpl Infect Dis 11: 257–265.

37. Mathers AJ, Stoesser N, Sheppard AE, Pankhurst L, Giess A, Yeh AJ, Didelot X, Turner SD, Sebra R, Kasarskis A, Peto T, Crook D, Sifri CD. 2015. Klebsiella pneumoniae carbapenemase (KPC)-producing *K. pneumoniae* at a single institution: insights into endemicity from whole-genome sequencing. Antimicrob Agents Chemother 59:1656–1663.

38. Enfield KB, Huq NN, Gosseling MF, Low DJ, Hazen KC, Toney DM, Slitt G, Zapata HJ, Cox HL, Lewis JD, Kundzins JR, Mathers AJ, Sifri CD. 2014. Control of simultaneous outbreaks of carbapenemase-producing enterobacteriaceae and extensively drug-resistant Acinetobacter baumannii infection in an intensive care unit using interventions promoted in the Centers for Disease Control and Prevention 2012 carbapenemase-resistant Enterobacteriaceae Toolkit. Infect Control Hosp Epidemiol 35: 810–817.

39. Mathers AJ, Carroll J, Sifri CD, Hazen KC. 2013. Modified Hodge Test versus Indirect Carbapenemase Test: Prospective Evaluation of a Phenotypic Assay for Detection of Klebsiella pneumoniae Carbapenemase (KPC) in Enterobacteriaceae. J Clin Microbiol 51: 1291–1293.

40. Mathers AJ, Poulter M, Dirks D, Carroll J, Sifri CD, Hazen KC. 2014. Clinical Microbiology Costs for Methods of Active Surveillance for Klebsiella pneumoniae Carbapenemase-Producing Enterobacteriaceae. Infect Control Hosp Epidemiol 35: 350–355.

41. CDC. 2009. Guidance for control of infections with carbapenem-resistant or carbapenemase-producing Enterobacteriaceae in acute care facilities. MMWR Morb Mortal Wkly Rep 58: 256–260.

42. Chen L, Chavda KD, Al Laham N, Melano RG, Jacobs MR, Bonomo RA, Kreiswirth BN. 2013. Complete nucleotide sequence of a *bla*_KPC_-harboring Incl2 plasmid and its dissemination in New Jersey and New York hospitals: a hidden threat. Antimicrob Agents Chemother 57: 6.

43. Kitchel B, Sundin DR, Patel JB. 2009. Regional dissemination of KPC-producing Klebsiella pneumoniae. Antimicrob Agents Chemother 53: 4511–4513.

44. Partridge SR, Hall RM. 2005. Evolution of transposons containing blaTEM genes. Antimicrob Agents Chemother 49: 1267–1268.

45. Bailey JK, Pinyon JL, Anantham S, Hall RM. 2011. Distribution of the blaTEM gene and blaTEM-containing transposons in commensal Escherichia coli. J Antimicrob Chemother 66: 745–751.

46. Zong Z, Ginn AN, Dobiasova H, Iredell JR, Partridge SR. 2015. Different IncI1 plasmids from Escherichia coli carry ISEcp1-blaCTX-M-15 associated with different Tn*2*-derived elements. Plasmid.

47. Rice LB, Carias LL, Hutton RA, Rudin SD, Endimiani A, Bonomo RA. 2008. The KQ element, a complex genetic region conferring transferable resistance to carbapenems, aminoglycosides, and fluoroquinolones in *Klebsiella pneumoniae*. Antimicrob Agents Chemother 52: 3427–3429.

48. Martínez T, Vázquez GJ, Aquino EE, Martínez I, Robledo IE. 2014. ISEcp1-mediated transposition of *bla*_KPC_ into the chromosome of a clinical isolate of Acinetobacter baumannii from Puerto Rico. J Med Microbiol 63: 1644–1648.

49. Mathers AJ, Cox HL, Kitchel B, Bonatti H, Brassinga AK, Carroll J, Scheld WM, Hazen KC, Sifri CD. 2011. Molecular dissection of an outbreak of carbapenem-resistant enterobacteriaceae reveals Intergenus KPC carbapenemase transmission through a promiscuous plasmid. MBio 2:e00204–00211.

50. Guindon S, Gascuel O. 2003. A simple, fast, and accurate algorithm to estimate large phylogenies by maximum likelihood. Syst Biol 52: 696–704.

51. Reeves PR, Liu B, Zhou Z, Li D, Guo D, Ren Y, Clabots C, Lan R, Johnson JR, Wang L. 2011. Rates of mutation and host transmission for an Escherichia coli clone over 3 years. PLoS One 6:e26907.

